# Confidence boosts serial dependence in orientation estimation

**DOI:** 10.1101/369140

**Authors:** Jason Samaha, Missy Switzky, Bradley R. Postle

**Affiliations:** University of California, Santa Cruz, Department of Psychology; University of Wisconsin-Madison, Department of Psychology; University of Wisconsin-Madison, Department of Psychiatry

**Author notes:** Correspondence: Jason Samaha, Psychology Faculty Services, University of California, Santa Cruz, 1156 High Street, Santa Cruz, CA 95064.

**Keywords:** Confidence, Serial dependence, Decision-making, Metacognition, Population code

## Abstract

In the absence of external feedback, a decision maker must rely on a subjective estimate of their decision accuracy in order to appropriately guide behavior. Normative models of perceptual decision making relate subjective estimates of internal signal quality (e.g. confidence) directly to the internal signal quality itself, thereby making it unknowable whether the subjective estimate or the underlying signal is what drives behavior. We constructed stimuli that dissociated human observer’s performance on a visual estimation task from their subjective estimates of confidence in their performance, thus violating normative principles. To understand whether confidence influences future decision making, we examined serial dependence in observer’s responses, a phenomenon whereby the estimate of a stimulus on the current trial can be biased towards the stimulus from the previous trial. We found that when decisions were made with high confidence, they conferred stronger biases upon the following trial, suggesting that confidence may enhance serial dependence. Critically, this finding was true also when confidence was experimentally dissociated from task performance, indicating that subjective confidence, independent of signal quality, can amplify serial dependence. These findings demonstrate an effect of confidence on future behavior, independent of task performance, and suggest that perceptual decisions incorporate recent history in an uncertainty-weighted manner, but where the uncertainty carried forward is a subjectively estimated and possibly suboptimal readout of objective sensory uncertainty.

## Introduction

Humans are capable of estimating the accuracy of their decisions even in the absence of external feedback. For example, subjective confidence ratings correlate with objective accuracy across a variety of perceptual and mnemonic tasks (Fleming et al., 2010; Song et al., 2011; Ais et al., 2016; Samaha and Postle, 2017), indicating that confidence depends, at least in part, on the same information underlying choices. This metacognitive ability may be crucial for adaptive behavior as it provides an estimate of performance that could be utilized in future decision processes such as optimizing decision policies (van den Berg et al., 2016), learning from mistakes (Yeung and Summerfield, 2012), or deciding to seek out new information (Call and Carpenter, 2001; Kepecs et al., 2008; Hayden et al., 2011).

Because confidence is correlated with task performance, however, it is difficult to know if subjective confidence *per se* influences subsequent behavior, or if the underlying sensory uncertainty on which confidence is based is sufficient to drive future behavior. Indeed, normative models of perceptual decision making posit a direct relationship between sensory uncertainty and the readout of subjective confidence (Kiani and Shadlen, 2009; Meyniel et al., 2015; Pouget et al., 2016; Sanders et al., 2016). Typically, experimenters manipulate stimulus evidence and evaluate the relation between decision accuracy and confidence (Kiani et al., 2014; van den Berg et al., 2016; Zylberberg et al., 2016). Or, stimulus evidence is kept constant and trial-to-trial covariation in confidence and accuracy is examined (Hebart et al., 2014). Both approaches, however, do not allow one to separate the influence of confidence from the influence of the quality of evidence. Either by manipulating evidence externally, as in the former case, or by relying on internal fluctuations of stimulus evidence, as in the latter case. Previous paradigms have not teased apart sensory uncertainty and subjective confidence when examining the effects of confidence on subsequent behavior (for critical review, see Samaha, 2015).

One exception is a recent study that employed two stimulus conditions that were equated in terms of accuracy (and hence evidence quality), but which differed in terms of confidence. By using these stimuli in a perceptual discrimination task that allowed subjects to collect additional evidence when they felt unconfident, the researchers showed that subjective confidence biased information seeking behavior even when accuracy was matched (Desender et al., 2018). Here, we apply the same logic to investigate whether subjective confidence, independent of the quality of evidence, modulates the influence of a current perceptual state on subsequent perceptual decisions, a phenomenon known as serial dependence (Fischer and Whitney, 2014).

Serial dependence often manifests as a bias towards reporting that a current stimulus appears more similar to recently seen stimuli than it actually is. Serial dependence occurs for a range of stimulus features, including luminance (Fründ et al., 2014), orientation (Fischer and Whitney, 2014; Fritsche et al., 2017), spatial location (Bliss et al., 2017), direction of motion (Alais et al., 2017), numerosity (Fornaciai and Park, 2018), motion variance (Suárez-Pinilla et al., 2018), and higher-level features such as face identity (Liberman et al., 2014). Although suboptimal in a psychophysical task where stimuli are temporally uncorrelated, in many real-world scenarios stimuli are stable across various time scales and serial dependence may be an adaptive bias that promotes temporal continuity (Kiyonaga et al., 2017). It was recently suggested that the influence of previous trials is mediated by observer’s confidence on those trials. Braun and colleagues found that the magnitude of history biases increased when responses on the previous trials were correct and faster, two proxies for confidence (Braun et al., 2018). This study, however, did not explicitly measure confidence, and, by design, the proxies for confidence that were used (accuracy and RT) are directly related to the quality of evidence. Suárez-Pinilla and colleagues also found that confidence on the previous trial modulated serial dependence in motion variance estimates, but also did not dissociate confidence from task performance (Suárez-Pinilla et al., 2018). Therefore, it is still unknown whether subjective confidence is capable of boosting serial dependence even when divorced from the quality of evidence (see Figure 1A).

**Figure 1.**
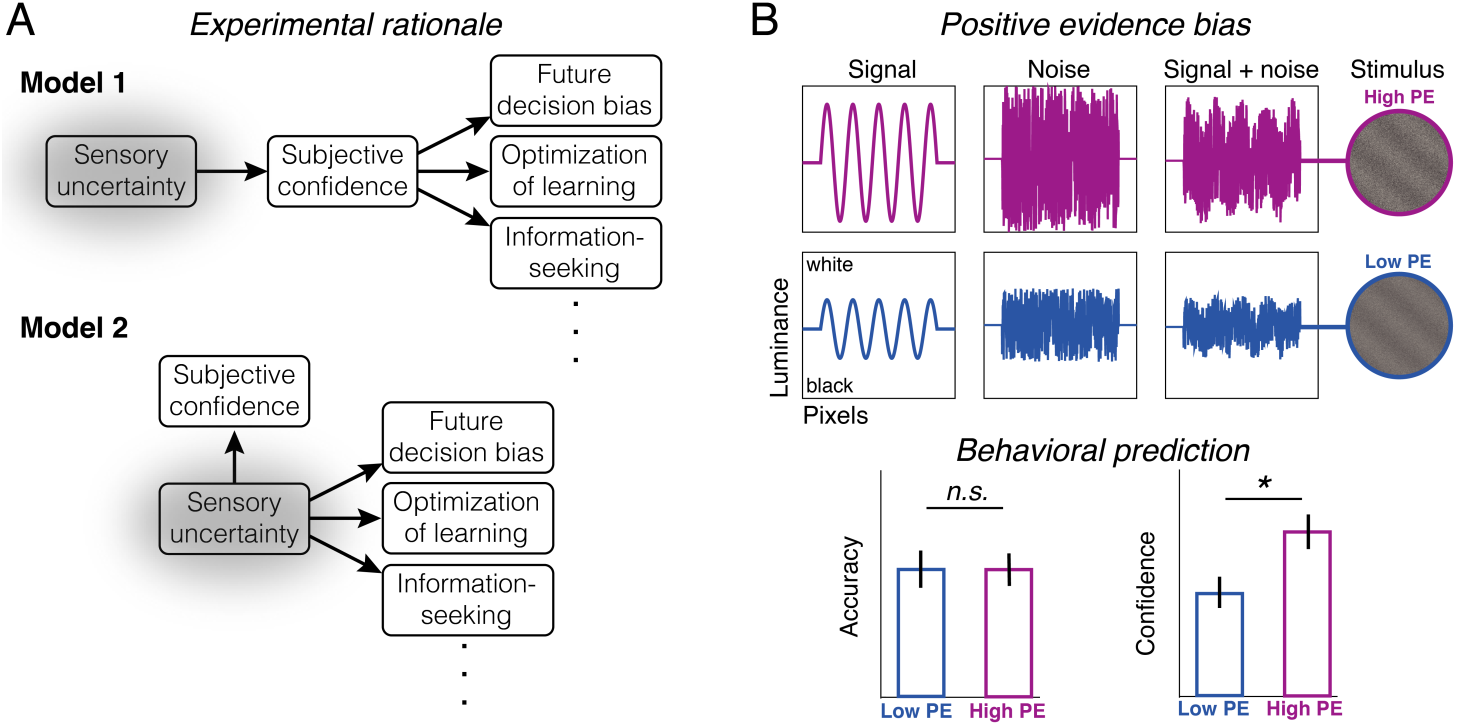
Experimental rationale, stimulus construction, and behavioral predictions. **A,** Subjective confidence informs future behavior by providing an estimate of sensory uncertainty. Most experimental evidence to date, however, is compatible both with a model in which subjective confidence directly informs future behaviors based on an estimate of sensory uncertainty (model 1) and with a model in which subjective confidence is epiphenomenal, but correlated with sensory uncertainty, and sensory uncertainty alone suffices to drive future behaviors (model 2). **B,** Teasing apart these models requires dissociating confidence from sensory uncertainty. We presented observers with sinusoidal luminance gratings averaged with white noise (upper panel). In the high positive evidence (PE) condition, stimuli had relatively high contrast noise and high contrast signal. In the low PE condition, signal and noise contrast was half of that in the high PE condition. Here, the term PE refers to the amount of contrast supporting correct stimulus identification (i.e., the amount of signal contrast). On the basis of prior work (Koizumi et al., 2015; Samaha et al., 2016), we predicted that the low PE condition would result in a decrease in confidence without changing the accuracy of orientation estimates (lower panel), a phenomenon we term the positive evidence bias (PEB).

Here, we capitalize on recent findings demonstrating that confidence judgments are overly reliant on the magnitude of evidence in favor of a perceptual decision, whereas decision accuracy is determined by the balance of evidence for each alternative (Zylberberg et al., 2012; Koizumi et al., 2015; Maniscalco et al., 2016; Peters et al., 2017; Rausch et al., 2017; Samaha et al., 2017; Odegaard et al., 2018). We recently showed that this can lead to a dissociation of confidence and accuracy by proportionally increasing the strength (in terms of visual contrast) of the signal and noise components of a grating + white noise stimulus during an orientation discrimination task (Samaha et al., 2016). This procedure effectively leaves the quality of evidence unchanged (thus, task performance is also unchanged). However, because positive evidence (i.e., the contrast of the grating component) is increased, this leads to increased confidence. We refer to this phenomenon as the positive evidence bias (PEB; where “positive evidence” refers to the amount of evidence in the stimulus supporting correct stimulus identification). Work so far, however, has demonstrated the PEB only in the context of discrimination tasks, where choice and confidence computations may differ from those employed in the continuous estimation tasks often used to demonstrate serial dependence (Fischer and Whitney, 2014; Liberman et al., 2014; Bliss et al., 2017; Fritsche et al., 2017; see *Discussion*).

The motivation for the present experiment is two-fold. First, we examined whether stimuli judged with higher confidence would produce larger biases on subsequent trials even when equating for task accuracy via the PEB. Second, we sought to replicate the PEB using a continuous orientation estimation task with confidence ratings, demonstrating the generality of the effect from Samaha et al., (2016).

## Materials and Methods

### Participants

20 participants were recruited from the University of Wisconsin-Madison (mean age = 20.6 years, *SD* = 2.01, 14 female). All subjects reported normal or corrected visual acuity, provided written informed consent, and were compensated monetarily. Sample size was chosen to be on par with recent serial dependence experiments which focus on group-level statistical inferences (Alais et al., 2017; Bliss et al., 2017; Fritsche et al., 2017), while also being large enough to detect the PEB, as per our prior work (Samaha et al., 2016). Data from this experiment were published previously as part of a multi-experiment study addressing different hypotheses (Samaha and Postle, 2017). This experiment was conducted in accordance with the University of Wisconsin Institutional Review Board and the Declaration of Helsinki. In accordance with the practices of open science and reproducibility, all raw data and code used in the present analyses are freely available through the Open Science Framework (https://osf.io/py38c/).

### Stimuli

Visual stimuli were composed of a sinusoidal luminance grating (1.5 CPD, zero phase) embedded in white noise and presented centrally within a circular aperture (2 DVA). The orientation of the grating was randomly chosen on each trial from the range 0:179° in integer steps. The noise component of the stimulus was created anew on each trial by randomly sampling each pixel’s luminance from a uniform distribution. The probe grating was rendered without noise at 30% Michelson contrast and was initiated at a random orientation on every trial to avoid response preparation. A fixation point (light gray, 0.08 DVA) was centered on the screen and was dimmed slightly to indicate trial onset (see Figure 2A). Stimuli were presented atop a gray background on an iMac computer screen (52 cm wide × 32.5 cm tall; 1920 × 1200 resolution; 60 Hz refresh rate) using the MGL toolbox (http://gru.stanford.edu) running in MATLAB 2015b (MathWorks, Natick, MA, USA) viewed from a chin rest at a distance of 62 cm.

**Figure 2.**
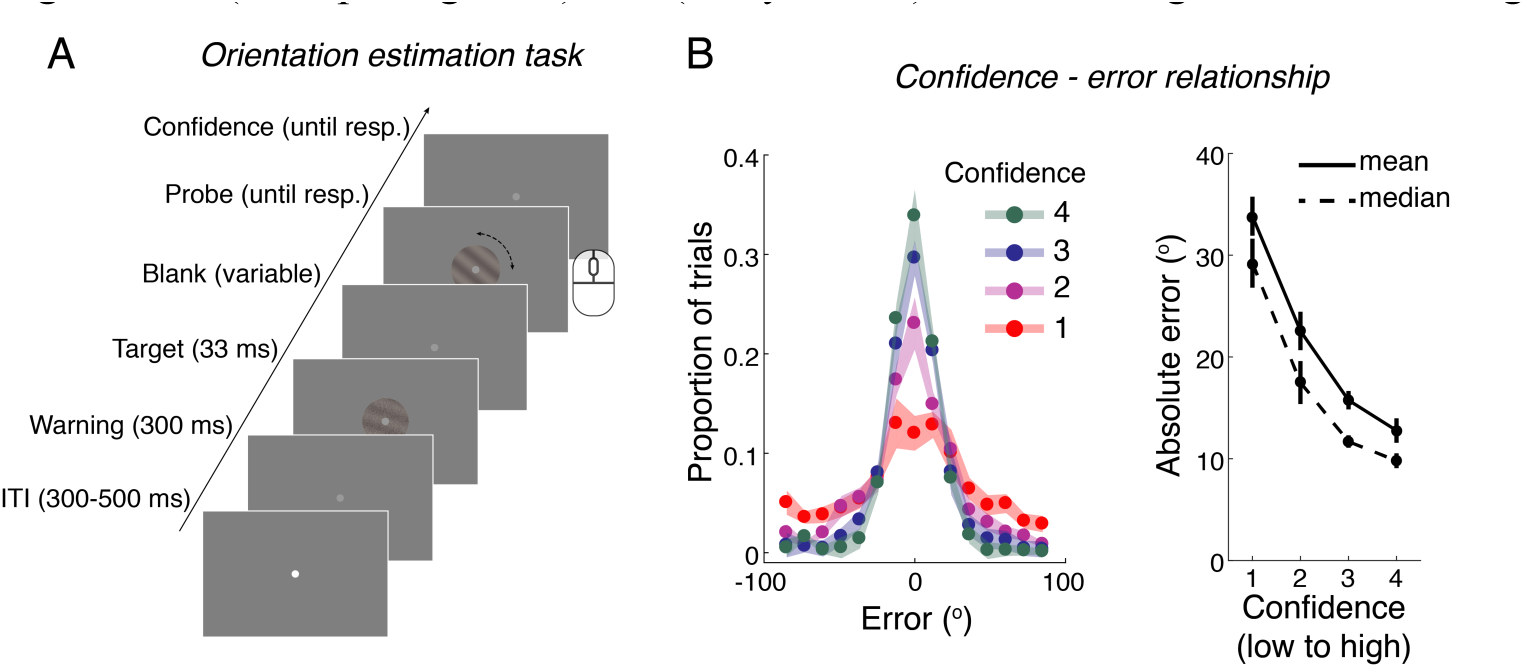
Task timing and confidence-error relationship. **A,** A target grating was briefly presented with a randomly determined orientation on each trial. Following a variable delay, a noiseless probe grating appeared and subjects used a computer mouse to rotate the probe until it matched the orientation of the target. A subsequent confidence rating was given on a 4-point numerical scale. Grating stimuli contained either high or low PE, randomly determined on each trial. **B,** The left panel shows the distributions of response errors as a function of confidence ratings. The right panel shown mean and median absolute error at each confidence level. Both plots reveal that error decreases with increasing confidence, suggesting that subjects have knowledge of the accuracy of their own orientation estimates and were generally using the confidence scale appropriately. Shaded bands and error bars denote ±1 SEM.

### Procedure

The subject’s task was to rotate a probe grating with a computer mouse to match the orientation of the target grating and then provide a confidence judgment. Subjects pressed the spacebar key to lock-in their orientation response and then used number keys 1-4 to rate a confidence. Because performance in this task varies continuously (as opposed to binary correct/incorrect outcomes), we instructed subjects to use the confidence scale to indicate how close they think they came to the true orientation using labels 1 (“complete guess”) to 4 (“very close”). Event timings are shown in Figure 2A.

Whereas previous experiments examining serial dependence for orientation have used grating stimuli well above contrast thresholds (Fischer and Whitney, 2014; Fritsche et al., 2017), we required stimuli to be near-threshold to replicate the PEB from prior work (Koizumi et al., 2015; Maniscalco et al., 2016; Samaha et al., 2016) and to ensure that the entire range of the confidence scale was used by subjects. We therefore began each experimental session with 100 trials of a 1-up, 3-down adaptive contrast staircase. To adapt the staircase to an estimation task, responses were classified as correct or incorrect depending on whether they were within ±25° of the true orientation. This procedure aimed to produce ∼80% of trials with less than ±25° error. The staircase began with the grating component of the stimulus having a Michelson contrast of 50%, which was then averaged with a 100% contrast white noise patch. The step size in grating contrast was adapted according to the PEST algorithm (Taylor and Creelman, 1967), with an initial starting step size of 20% contrast. The resulting mean contrast of the grating (prior to averaging with 100% noise) was 8.5% (*SD* = 2.72), which was held constant throughout the subsequent main task.

For the main task, we presented stimuli from two conditions: a high positive (PE) condition and a low positive evidence condition. Following our prior work (Samaha et al., 2016), the contrast of stimuli in the high PE condition were taken directly from the staircase procedure^1^, whereas the contrast of the grating and the noise component of the stimuli in the low PE condition were both halved with respect to the high PE values (see Figure 1B). In other words:

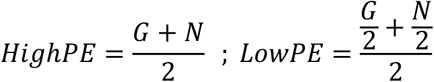

where G is the contrast of the grating component of the stimulus defined from the staircase, N is the contrast of the noise component (which was set to100%), and *HighPE* and *LowPE* refer to the stimuli in the high and low PE conditions, respectively. This procedure matches the signal-to-noise ratio across both conditions, which we anticipate would lead to no change in estimation accuracy, but would lead to a change in confidence, if confidence is over-reliant on the magnitude of G, a proxy for the amount of PE represented in the brain.

A high or low PE stimulus was chosen randomly for each trial. As recent work has suggested that serial dependence becomes stronger when the target stimulus is held in short-term memory (Bliss et al., 2017; Fritsche et al., 2017), we randomly sampled the duration of the delay between the stimulus and the probe grating from the following values (in seconds): 0.6, 3.45, 6.3, 9.15, and 12. Subjects completed 300 trials of the main experiment, divided evenly into 5 blocks. Total task time was approximately 1.5 hours.

### PEB analysis

Error was computed for each trial as the angular distance between the target orientation and the response (Figure 3A). We quantified accuracy on high and low PE trials using four metrics: The median and mean of the absolute response error as well as the precision and guess rate obtained from a two-component mixture model fit to the distribution of response errors for each subject (Bays et al., 2009). The latter two metrics are obtained via fitting a mixture of a Von Mises and a uniform distribution to response errors, resulting in a concentration parameter, *κ*, which describes the precision of the Von Mises, and a parameter that describes the height of the uniform distribution, which corresponds to the probability of making a random (“guess”) response. The model was fit to data using an expectation-maximization algorithm implemented in MATLAB code obtained from www.bayslab.com. We did not obtain enough trials at each of the five delay durations to reliably fit mixture models to each combination of delay and PE level separately. Therefore, any analysis of PEB with delay as a factor was conducted on mean and median absolute error. Confidence for high and low PE conditions was quantified as the mean rating across each type of trial. The effect of PE on confidence and each accuracy metric was evaluated statistically using two-tailed paired-sample t-tests. Improbable trials with responses faster than 200 ms or slower than the 95th percentile of response times across all subjects (5.04 sec) were discarded prior to any analysis. Lastly, because we predict a null effect of our PE manipulation on estimation accuracy, we include Bayes factors whenever interpreting null hypotheses. All Bayes factors are reported in terms of evidence for the null hypothesis (BF_null_), quantified as how many times more likely the data are to be observed under the null hypothesis. In the case of t-tests, JZS BFs were computed according to Rouder et al. (2009) using a normal prior (Jarosz and Wiley, 2014). In the case of correlations BFs were computed according to (Wetzels and Wagenmakers, 2012).

**Figure 3.**
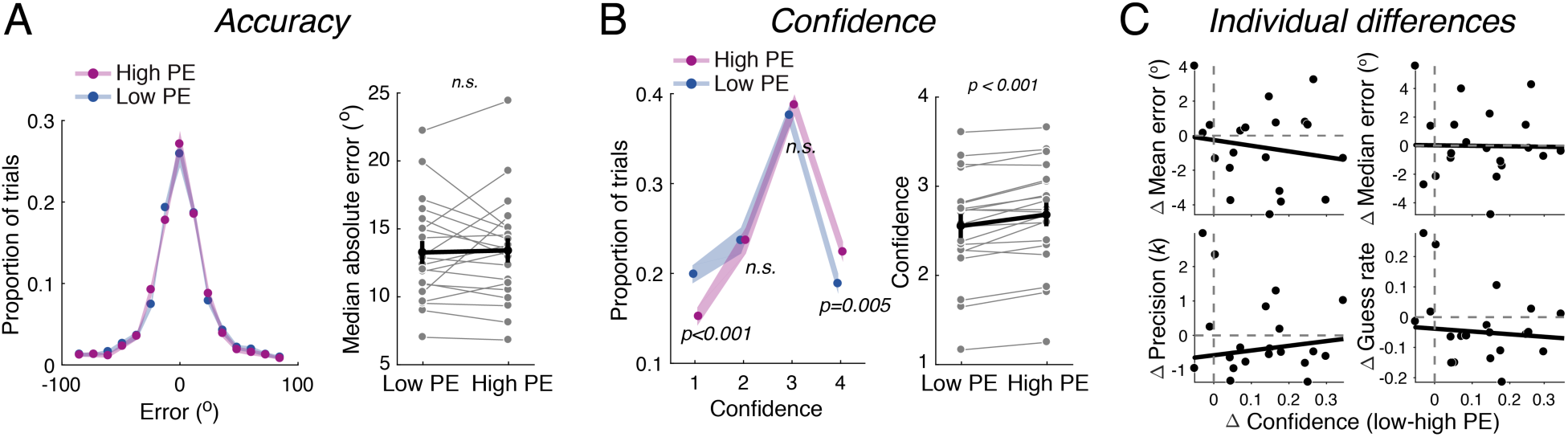
The positive evidence bias (PEB) in orientation estimation. **A,** Left panel depicts the distribution of response errors for high and low PE stimuli, binned and averaged across subjects. Right panel shows median absolute response error for each subject. Shaded bands and error bars denote ±1 SEM. The overlap of error distributions and lack of reliable accuracy changes suggests that PE levels did not noticeably impact estimation accuracy (see Results for additional quantifications of accuracy). **B,** Left panel shows the distribution of responses at each level of confidence as a function of PE. Increasing PE lead to a significant increase in highly confident responses (“4”) and a decrease in low confident (guessing) responses (“1”). Mean confidence (right panel) was higher for high PE stimuli, a bias present in 17/20 subjects. Error bars span ±1 SEM, shaded bands cover ±1 within-subject 95% CI (Morey, 2008) **C,** Correlations between individual differences in PE-related variability in confidence (x-axis in all plots) and PE-related variability in accuracy across four accuracy metrics (subtraction is always low-high PE). Lines denote robust linear fit. No correlations were significant. Collectively, these results suggest our stimulus manipulation selectively modulated confidence without changing accuracy.

### Serial dependence analysis

Several preprocessing steps were taken prior to estimating the magnitude of serial dependence. Following others (Bliss et al., 2017; Fritsche et al., 2017), trials with high error were discarded. Since we intentionally staircased performance by classifying trials as correct if they were within ± 25° error, we applied this same threshold to remove incorrect trials prior to quantifying serial dependence. This step ensured that trials that were likely unperceived were not included in the analysis. Indeed, this step was necessary to observe any reliable serial dependence at all (see Results). Next, response errors were demeaned by subtracting each subject’s mean (signed) error from the error on each trial. By subtracting each subject’s average error from each trial, this step removes any small clockwise or counter-clockwise response biases (Bliss et al., 2017; Fritsche et al., 2017).

We quantified serial dependence using three methods: a model-based, a model-free, and a Fourier-based analysis. For the model-based analysis, we sorted error on the current trial by the relative difference in orientation between the stimulus on the previous and current trial (see Figure 4). The first trial of each block was not considered to have a previous trial. If a trial was removed due to inaccuracy, then the most immediately preceding correct trial was considered the “previous trial” (this is not unreasonable, as serial biases have been shown to extend at least 3 trial back (Fischer and Whitney, 2014). If orientation responses are biased towards the previous trial then error on the current trial (y-axis) will be pulled towards the same sign as the relative difference (x-axis), whereas a repulsive bias would result in response errors of an opposite sign, and no serial dependence would result in a flat line. This profile has been previously parameterized by fitting the data with a derivative-of-Gaussian function (DoG; Fischer and Whitney, 2014; Liberman et al., 2014; Alais et al., 2017; Bliss et al., 2017; Fritsche et al., 2017) of the form:

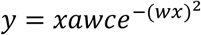

where *x* is the relative orientation of the previous trial, *a* is the amplitude of the curve peaks, *w* is the width of the curve and *c* is the constant 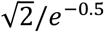, which scales the amplitude parameter of interest to numerically match the height of the curve in degrees. Following others (Bliss et al., 2017; Fritsche et al., 2017), we fit this function to group-averaged data after first smoothing individual subject’s data with a 25-trial moving median filter (changes in filter size within a reasonable range did not change the results). The amplitude parameter *a* and width parameter *w* were free to vary across a wide range of plausible values between [-15°, 15°] and [0.02, 0.2] respectively. Fitting was conducted by minimizing the sum of squared errors using the MATLAB routine *lsqcurvefit.m*. To determine the statistical significance of group-level DoG fits, we used a bootstrapping procedure (DiCiccio and Efron, 1996). On each of 80,000 iterations we sampled subjects with replacement and fit a DoG to the average of the bootstrap sampled data. We saved the value of the amplitude parameter after each iteration, forming a distribution of the amplitude parameter of our sample. We computed 95% confidence intervals from this distribution and a p-value was calculated as the proportion of samples above zero amplitude (no serial dependence), which was considered significant at α = 0.025 (two-tailed bootstrap test). To test whether confidence or PE on the previous trial predicted serial biases on the current trial, we refit DoG functions to data split according to whether the previous trial was high or low confidence (mean split according to each subject’s mean confidence rating), or high or low PE. Statistical significance testing was conducted using the same bootstrap procedure as above, but the difference in serial dependence amplitude between conditions was saved on each iteration and a p-value was computed as the proportion of the difference score distribution greater than zero (α = 0.025, two-tailed bootstrap test).

**Figure 4.**
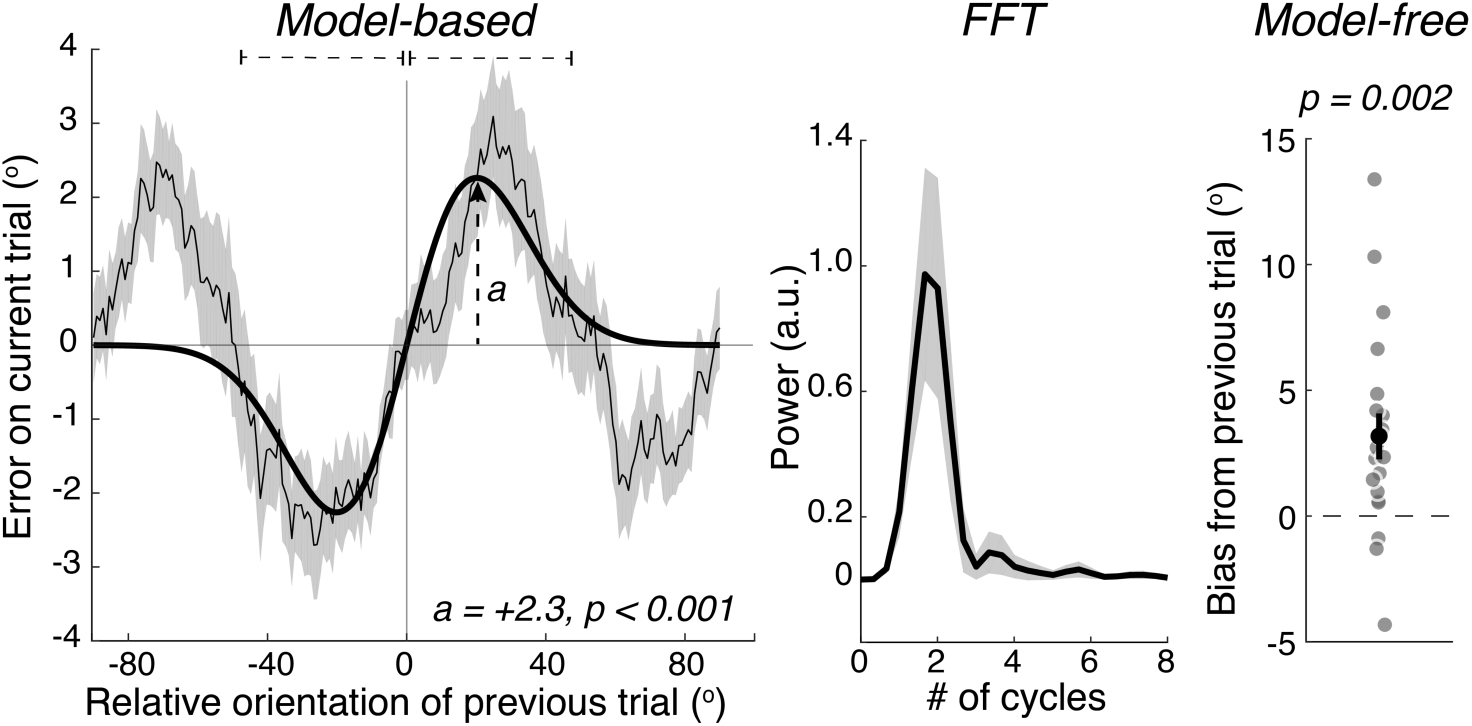
Serial dependence in orientation estimates. Left panel shows error on the current trial as a function of the difference between the orientation on the current and previous trial. The thick black line shows the fit of a DoG model to the smoothed group-level data. Negative values indicate counterclockwise differences. The amplitude parameter, a, of the DoG function reflects the height of the function and captures the magnitude of bias due to the orientation on the previous trial. Positive/negative a denotes an attractive/repulsive bias. In keeping with other results, close relative orientations (within 45°) lead to a significant attractive bias which turns to a repulsive bias when the relative orientation on the previous trial was further in stimulus space (greater than 45°). Note that the DoG captures only the attractive bias at smaller relative differences. Dashed lines above the plot denote the windows used to estimate serial bias in the model free analysis (right panel), which also revealed reliable serial dependence. The middle panel depicts the power spectrum of the curve in the left plot. This reveals a clear periodicity at around 2 cycles, reflecting the sinusoidal nature of the serial dependence profile and better capturing the repulsive bias at larger relative orientation differences. The amplitude at this peak frequency serves as another quantification of the magnitude of serial dependence using a basis set of sinusoids. Shaded bands and error bars in all figures span ±1 SEM.

To ensure that our results were not a quirk of model fitting, we additionally tested for serial dependence and its modulation by confidence using a model-free analysis. For each subject, we computed the median (signed) error across trials where the relative difference between the current and previous stimulus fell within the interval (0°, 45 °], and subtracted that from the median error on trials within the interval [-45 °, 0) (see dashed lines in Figure 4). Thus, positive and negative values indicate an attractive or repulsive bias, respectively. This metric was computed for all levels of confidence and PE on the previous trial. The influence of delay on the previous trial was also tested this way, as there were insufficient trial numbers at each delay to fit with a DoG. Statistical testing was performed using two-tailed paired-samples t-tests, or by fitting a linear function to each subject’s bias by confidence or bias by delay data and comparing the slope against zero at the group level with a two-tailed paired-samples t-tests (Figure 5A, right panel).

**Figure 5.**
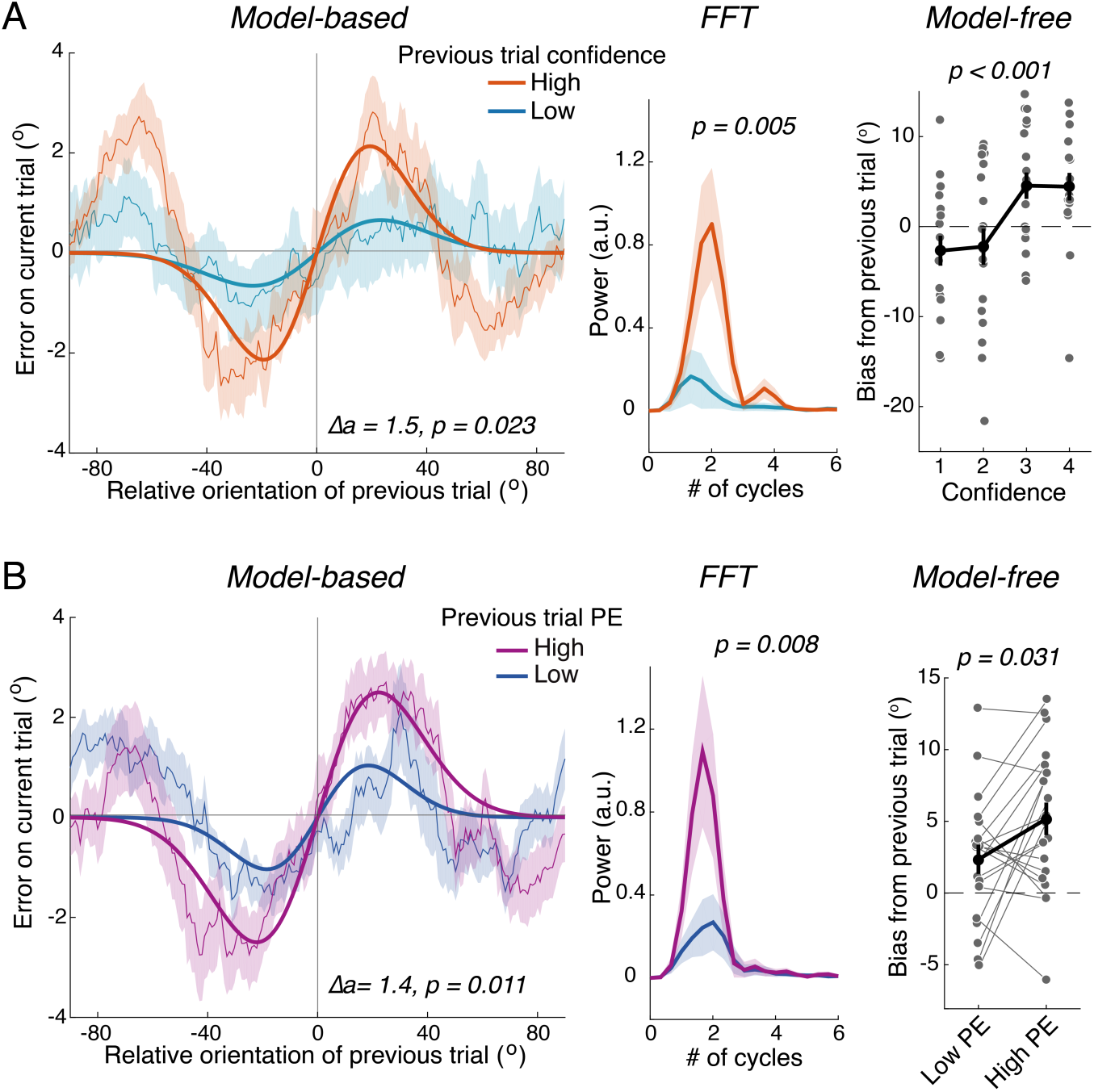
Confidence boosts serial dependence. **A,** Left panel shows serial dependence curves and DoG fits to data separated according to whether confidence on the previous trial was high or low. High confidence on the previous trial was associated with increased serial dependence amplitude in the DoG model-based analysis and the corresponding FFT analysis of the serial dependence curves (middle panel). The model-free analysis at each level of confidence (right panel) also showed that serial biases increased with increasing confidence on the previous trial. **B,** Sorting data by PE on the previous trial revealed that trials with high PE more strongly biased responses on the subsequent trial in all three analyses: The DoG model-based, the FFT, and model-free analysis. This suggests that increasing confidence without changing accuracy is sufficient to boost serial biases. Shaded bands and error bars are ±1 SEM. Δa refers to the difference in amplitude parameter between conditions.

The serial dependence curves revealed an additional repulsive bias at larger orientation differences that was not captured well by the DoG fit used in prior literature (Figures 4 and 5; see also Bliss et al., 2017; Fritsche et al., 2017). Because these bumps in the serial dependence profile essentially make the curves sinusoidal, we decomposed these curves into sine waves of varying amplitude, phase, and frequency using a fast Fourier transform (FFT). The group-level curves shown in Figures 4 and 5 were zero padded (frequency resolution 0.33 Hz), linearly detrended, and then transformed into power spectra by squaring the absolute value of the complex FFT result (MATLAB function *fft.m*). Serial dependence was quantified as the power at the frequency with highest power for each condition (the “dominant frequency”). This is akin to finding the amplitude of the best-fitting sinusoid that is allowed to vary in both frequency and phase. Following the statistical analysis of the DoG fit (described above), a bootstrap procedure was performed whereby group-level serial dependence curves were recomputed using a random subset of subjects (sampled with replacement) and decomposed with an FFT. On each of 80,000 bootstrap iterations, the power spectrum for each condition (Figure 5) was saved and the power of the dominant frequency in each condition was recorded. Subtracting the distributions created a distribution of difference scores reflecting the change in sine wave amplitude across conditions. Using these difference-score distributions, we computed p-values by taking the proportion of bootstrap samples greater than 0 (α = 0.025, two-tailed bootstrap test) as well as 95% confidence intervals. Note that the x-axis in the FFT plots are normalized to reflect the # of cycles of a particular sine wave across the whole serial dependence plot, rather than Hz (since the data are not a timeseries).

## Results

### PEB for orientation estimation

We analyzed accuracy and confidence ratings during an orientation estimation task where stimuli contained either high or low PE, but were matched for overall signal-to-noise ratio. Across all trials, estimation error sharply decreased with increasing confidence (Figure 2B) and single-trial Spearman correlations between absolute error and confidence revealed negative relationships for every participant (rho range: [-0.46 -0.01]). This indicates that subject’s confidence ratings reflected some knowledge of their own performance. A repeated measures ANOVA including an interaction term between delay duration and PE did not reveal any reliable interaction of PE with delay when predicting accuracy (mean or median absolute error) or confidence (all *ps* > 0.05), therefore we focus on paired comparisons between high and low PE trials, aggregating over delay duration. As hypothesized, proportionally increasing both signal and noise contrast in a compound grating stimulus lead to no discernable change in the accuracy of observer’s responses as characterized by median response error (*t*(19) = -0.27, *p* = 0.79, BF_null_=5.66), mean response error (*t*(19) = 1.20, *p* = 0.24, BF_null_=3.00), the precision of responses (see Materials and Methods; *t*(19) = 0.17, *p* = 0.86, BF_null_=5.78), or the probability of making a random response (*t*(19) = 1.10, *p* = 0.28, BF_null_=3.33). See Figure 3A,C. This is in line with previous null effects of the exact same (Samaha et al., 2016) and similar PE manipulations on 2-choice discrimination accuracy (Zylberberg et al., 2012; Koizumi et al., 2015; Maniscalco et al., 2016). The Bayes factor analysis indicates that, across different metrics, the change in accuracy we observed is between 3 and 5.78 times more likely to be observed under the null.

In contrast to the null result of PE on accuracy, we observed a highly significant modulation of subjective confidence ratings, such that mean confidence was greater for high as compared to low PE stimuli (*t*(19) = -5.06, *p* = 0.00006; Figure 3B). Analysis of the proportion of responses at each of the four confidence levels (Figure 3B) revealed that increasing PE led a decrease in the use of “1” ratings (“complete guess”) and an increase in the use of “4” ratings (“very close to the true orientation”; *p-*value per level of confidence: *p*_conf1_ = 0.0002, *p*_conf2_ = 0.99, *p*_conf3_ = 0.35, *p*_conf4_ = 0.005). Additionally, we checked whether individual differences in the PE-related change in accuracy and the PE-related change in confidence were correlated. Across all four metrics of accuracy, there was virtually no correlation (Spearman’s rho) across subjects (mean error: *rho* = -0.073, *p* = 0.75, BF_null_=5.60; median error: *rho* = -0.024, *p* = 0.92, BF_null_=5.83; precision: *rho* = -0.066, *p* = 0.78, BF_null_=5.64; guess rate: *rho* = -0.176, *p* = 0.45, BF_null_=4.45; Figure 3C). This provides further evidence of independence between confidence and accuracy, indicating that even for an individual whose accuracy benefited from increasing PE, their confidence did not increase in kind. Analysis of Bayes factors suggest that the correlation between individual differences are 4.45 to 5.83 times as likely to be observed under the null hypothesis of no correlation. This result suggests that confidence is not simply a more sensitive measure of behavior than estimation error, as individual differences would likely be correlated under this hypothesis.

### Subjective confidence amplifies serial dependence

We characterized serial dependence by fitting a DoG function to group-level error expressed as a function of the relative orientation difference between the previous and current trial. As shown in Figure 4, trials showed a significant serial bias, such that responses were biased towards the orientation on the previous trial when the previous trial was within ∼ 45° of the current trial (serial dependence amplitude, *a* = 2.3°, 95% CI = [1.30 3.28], *p* < 0.0001), and a notable repulsive bias at larger relative orientation differences, consistent with recent reports (Bliss et al., 2017; Fritsche et al., 2017). This repulsive-attractive-attractive-repulsive profile produced an oscillation-like profile, which was verified with an FFT analysis showing a clear peak at a frequency of ∼2 cycles (Figure 4, middle panel). Serial dependence was undetectable when trials considered incorrect (see Methods) were included (*a* = 1.01°, CI = [-0.97 2.60], *p* = 0.11), which is sensible given that an undetected stimulus would not be expected to influence subsequent responses. The presence of serial dependence was also confirmed in the model-free analysis, which revealed a bias of comparable magnitude for trials within ±45° of relative difference (mean bias = 3.1°, CI = [1.26 5.07], *t*(19) = 3.45, *p* = 0.002). Serial bias showed no reliable linear relationship with delay duration on the previous (mean slope = -0.12, *t*(19) = -0.57, *p* = 0.57, BF_null_=5.02), or current trial (mean slope = 0.083, *t*(19) = 0.54, *p* = 0.59, BF_null_=5.10), so we focused subsequent analysis on serial dependence averaged across all delays.

Before addressing whether selectively increasing confidence by increasing PE caused an amplification of serially dependent biases, we first asked whether trial-to-trial variability in confidence predicted serial dependence. As shown in Figure 5A the DoG fit to trials preceded by high or low confidence responses revealed significant serial biases following high (*a* = 2.15°, CI = [1.25 3.04], *p* = 0.0001), but not low confidence trials (*a* = 0.64°, CI = [-0.78 1.82], *p* = 0.16). The difference distribution formed by subtracting the bootstrapped distribution of amplitude parameters in each condition was predominantly greater than zero, (Δ*a* = 1.50, CI = [0.003 3.21], *p* = 0.023), indicating significantly larger biases following high as compared to low confidence trials. The same conclusion was reached with an FFT analysis comparing power at the dominant frequency of serial dependence curves for each condition (Δpower = 0.73, CI = [0.17 1.35], *p* = 0.005; Figure 5A, middle panel). This effect was also confirmed in the model-free analysis: fitting a line to each subject’s serial bias magnitude as a function of their confidence on the previous trial revealed a significant positive relationship (mean slope = 2.98°, CI = [1.50 4.46], *t*(19) = 4.22, *p* = 0.0004). Paired contrasts at each level of confidence revealed that serial dependence was present only at confidence level 3 (*p* = 0.0073) and 4 (*p* = 0.020), and not levels 1 (*p* = 0.17) or 2 (*p* = 0.29; Figure 5A, right panel). These results suggest that confidence on the current trial may mediate that trial’s attractive influence on the subsequent trial. However, this finding conflates confidence with other factors that may relate to performance. For instance, if subjects were inattentive on the current trial, then confidence and performance could both be reduced, leading to a smaller bias on the subsequent trial. In this explanation, attention would be the primary variable leading to reduced serial bias, not subjective confidence.

Our task design teased apart confidence and performance by holding task performance constant while selectively increasing subjective confidence. We first checked that the PEB held for the subset of trials used for the serial dependence analysis. Indeed, across all four metrics of accuracy, there was no discernable difference according to the level of PE in the stimulus (all *ps* > 0.14, BF_nulls_ between 2.03 and 4.34). In fact, all metrics were pointing towards a difference in the opposite direction of confidence— slightly higher mean and median error, and lower *k* in the high PE condition (guess rate was 0 in all cases since high error trials were removed). Confidence, on the other hand, remained significantly higher for high PE stimuli (*t*(19) = -4.62, *p* = 0.0002), confirming the PEB for this subset of trials. DoG models fit to data sorted by high or low PE on the previous trial revealed significant serial dependence amplitudes following high PE trials (*a* = 2.52°, CI = [1.54 3.49], *p* = 0.00001), and an effect following low PE trials (*a* = 1.04°, CI = [0.12 1.84], *p* = 0.022). Critically, the distribution of amplitude differences from the bootstrap was significantly non-overlapping zero (Δ*a* = 1.48, CI = [0.21 2.90], *p* =0.011; Figure 5B, left panel), indicating that high PE on the previous trial lead to larger biases on the current trial than did low PE. The FFT analysis corroborated this effect, with higher power at the peak serial dependence frequency following high as compared to low PE trials (Δpower = 0.82, CI = [0.20 1.30], *p* = 0.008; Figure 5B, middle panel). The model-free analysis also replicated this result, with a significant bias following high (*t*(19) = 4.57, *p* = 0.0002) and low PE trials (*t*(19) = 2.23, *p* = 0.031), and a significantly greater bias following high as compared to low PE trials (*t*(19) = 2.32, *p* = 0.031; Figure 5B, right panel). Because high PE was associated with a boost in confidence, but no change in accuracy, these results suggest that increasing confidence independently of accuracy is sufficient for amplifying serial dependence in orientation judgments.

## Discussion

Many researchers have posited that the ability to assign confidence to one’s own performance serves a crucial role in formulating future behaviors (Yeung and Summerfield, 2012; Weil et al., 2013; Meyniel et al., 2015; van den Berg et al., 2016). The bulk of experimental work to date, however, has been unable to separate effects of subjective confidence from effects of task performance. For instance, a decision experienced with low confidence may alter future decision-making not because of the felt sense of confidence *per se*, but because attention on that trial was diverted and the stimulus was processed suboptimally. To ascertain whether subjective confidence can modulate dependencies between current and future decisions we designed an orientation estimation experiment that disentangled confidence ratings from objective task performance. We found that trial-to-trial variation in confidence predicted the magnitude of serial biases, such that when a trial was performed with high confidence it exerted a larger bias on the decision in the subsequent trial. Crucially, this relationship was replicated when we experimentally manipulated confidence levels without affecting task performance, indicating that confidence, divorced from performance, is capable of increasing serial dependence. This finding suggests that a representation of sensory uncertainty is carried forward to subsequent trials to influence decision-making. However, the representation of uncertainty that is carried forward need not be a perfect reflection of the actual stimulus evidence used to perform the task. Our results support a framework in which a suboptimal readout of sensory evidence forms the basis of subjective confidence judgments and gets carried forward to alter future decision behavior.

Why should confidence boost serial dependence? In the context of psychophysical experiments, serial biases are suboptimal because they lead to greater error when stimulus features are temporally uncorrelated. In real life, however, many stimuli are sufficiently auto-correlated (e.g., a book on a desk typically maintains some visual features from one second to the next) such that taking information from the recent past into consideration when making current decisions could be adaptive (Fischer and Whitney, 2014; Kiyonaga et al., 2017; Braun et al., 2018). As in other information integration problems, such as cue combination (Ernst and Banks, 2002), optimal integration of current and past sensory information requires weighting each representation by the uncertainty associated with it. In this way, recent sensory inputs can be thought of as a prior on current stimulus estimates (Bergen and Jehee, 2017). When the prior is associated with high uncertainty (low confidence) it should be given less weight in the current decision and thus lead to a smaller serial bias, as we observed. In this framework, though, our results suggest that the weights on the prior are determined not by the actual sensory uncertainty (which we equated) but by the biased readout of sensory uncertainty underlying subjective reports of confidence.

Biased estimates of confidence have been found to drive other decision-related behaviors as well. A recent experiment used a stimulus manipulation related to the one used here to manipulate confidence and accuracy independently (Desender et al., 2018). Consistent with our findings, the researchers also observed that selectively modulating confidence was sufficient to induce changes in future decision behavior, in the form of seeking additional information when confidence was low. Notably, though, two recent experiments have applied similar experimental manipulations of confidence and failed to find effects. Using the PEB in an orientation working memory task, we recently found no evidence that selectively modulating perceptual confidence led to changes in subsequent memory performance (Samaha et al., 2016). Furthermore, Koizumi et al (2015) used PEB-inducing stimuli as cues in a response inhibition task and in a response preparation task. Although they successfully increased confidence without changing performance, this change did not lead to enhanced performance in either task (Koizumi et al., 2015). Although there is little work using a dissociation paradigm such as the PEB to examine the function of subjective confidence, it is clear that not all tasks are affected by selectively modulating confidence. Such effects may be restricted to tasks involving an ongoing updating of decision policies or weighting of information in decision making (e.g., history biases, information-seeking, etc.).

The PEB is among a growing number of empirical demonstrations of a dissociation between objective task performance and subjective confidence (for review see Fleming and Daw (2017) and Rahnev and Denison (2018)). To our knowledge, however, such a dissociation has not yet been demonstrated in the context of a continuous estimation task, such as that used here. This is non-trivial because decision models based on continuous report performance often treat confidence as an optimal (in the sense of perfectly tracking accuracy) readout of sensory uncertainty (Meyniel et al., 2015). In In the framework of probabilistic population coding (Pouget et al., 2000; Ma et al., 2006; Beck et al., 2008), an ideal observer of neural activity could estimate the stimulus based on maximum likelihood (ML) decoding of the population activity and provide confidence by computing the width of the associated posterior distribution (the probability distribution of the stimulus conditioned on the observed spiking activity; Bays, 2016). This normative solution, however, fails to capture the PEB demonstrated here. Instead, we speculate that a neurally plausible computation of confidence based on the sum of activity across the population could account for the PEB in orientation estimation. Typically, the sum of activity across the population is inversely proportional to the width of the posterior and could therefore inform confidence (Ma et al., 2006; Meyniel et al., 2015; Bays, 2016). We reason that increasing the contrast of both signal and noise in our stimuli could lead to increased firing across all neurons in the population (Figure 6, right panel). This is plausible because responses in early visual cortex increase monotonically with contrast (Dean, 1981; Boynton et al., 1999) and because the stimulus manipulation used here is non-specific with respect to orientation contrast. If confidence is read out from this population via the sum of activity across it, confidence will be higher for our high PE stimuli whereas the ML estimate of the orientation will be unaffected. It is possible that divisive normalization (Heeger, 1992) may operate across this population, which could effectively cancel out this baseline increase in firing. We speculate, though, that this would only be true if the neurons in the normalization pool are exactly the same neurons used in the population read out of confidence, for example, if the sum of all neurons in the hypothetical population in Figure 6 is used in the denominator of the normalization equation (Carandini and Heeger, 2012).

**Figure 6.**
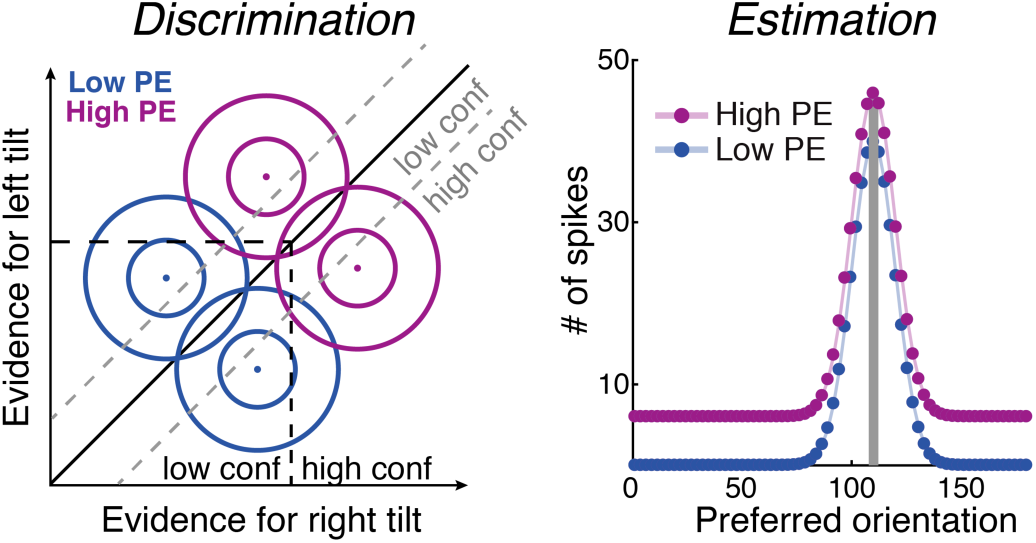
Possible decision models for the PEB. Left panel, in an orientation discrimination task modeled with signal detection theory the PEB could manifest if high/low confidence is rated via criteria placed along an axis perpendicular (black dotted lines) to the decision axis (diagonal black line), as opposed to parallel (gray dotted lines) to the decision axis (Maniscalco et al., 2016; Samaha et al., 2016). This corresponds to a confidence judgment based on the amount of evidence for a choice, rather than the balance of evidence for each choice. Circles denote two-dimensional Gaussian distributions corresponding to the internal sensory responses across trials for left or right orientation detectors. The PE manipulation is modeled as a translation of the distributions diagonally, without changing their separability (d’). Computations underlying confidence and accuracy in estimation tasks have been formulated in the framework of population coding. The right panel shows an idealized (noiseless) response across a population of neurons each tuned to different orientations for a high and low PE stimulus. The PEB could manifest if increasing signal and noise boosts the activity across all neurons in the population by an additive constant. Maximum likelihood decoding of the stimulus based on these population responses would result in the same orientation estimate (same peak value), but if confidence is read out via the sum of activity across the population, as suggested by prior work (Ma et al., 2006; Meyniel et al., 2015; Bays, 2016), then it would be higher for high PE stimuli.

In summary, we demonstrate a novel dissociation of confidence and performance in orientation estimates which has a possible neural grounding in current models of decision making, but which violates normative models of confidence. We show that orientation responses are serially dependent and that trials associated with high confidence, independent of task performance, confer larger biases upon subsequent trials. We interpret this finding as evidence that current decisions are biased by the recent past in a manner that is sensitive to the subjectively estimated uncertainty of recent inputs, thereby promoting uncertainty-weighted integration of current and future information.

## Footnotes

1 In Experiment 1 of Samaha & Postle (2017), a similar stimulus manipulation was used but under slightly different task conditions. In this experiment, halving signal and noise contrast led to a decrease in accuracy as well as confidence. For future work with this paradigm, we recommend individually staircasing both high and low PE stimuli to ensure that accuracy is matched, rather than hoping for matched accuracy after staircasing the high PE condition and halving the signal and noise contrasts to form the low PE condition, as was done here and in Samaha and Postle (2017).

